# ROCK1/Drp1-mediated aberrant mitochondrial fission is crucial for dopaminergic nerve cell apoptosis

**DOI:** 10.1101/366401

**Authors:** Qian Zhang, Changpeng Hu, Jingbin Huang, Wuyi Liu, Wenjing Lai, Faning Leng, Qin Tang, Yali Liu, Qing Wang, Min Zhou, Fangfang Sheng, Guobing Li, Rong Zhang

## Abstract

Dopamine deficiency caused by apoptosis of the dopaminergic nerve cells in the midbrain substantia nigra is the main pathological basis of Parkinson's disease (PD). Recent research has shown that dynamin-related protein 1 (Drp1)-mediated aberrant mitochondrial fission plays an important role in dopaminergic nerve cell apoptosis. However, the upstream regulatory mechanism remains unclear. Our study shows that knockdown of Drp1 blocked aberrant mitochondrial fission and dopaminergic nerve cell apoptosis. Importantly, we found that ROCK1 was activated in an MPP^+^-induced PD cell model and that ROCK1 knockdown and the specific ROCK1 activation inhibitor Y-27632 blocked Drp1-mediated aberrant mitochondrial fission and apoptosis of dopaminergic nerve cell through suppression of Drp1 dephosphorylation/activation. Our *in vivo* study confirmed that Y-27632 significantly improved symptoms of a PD mouse model through inhibition of Drp1-mediated aberrant mitochondrial fission and apoptosis of dopaminergic nerve cell. Collectively, Our study suggests an important molecular mechanism of PD pathogenesis involving ROCK1-regulated dopaminergic nerve cell apoptosis via activation of Drp1-induced aberrant mitochondrial fission.

## Introduction

Parkinson’s disease (PD), which often occurs in elderly patients, is a neurodegenerative disease characterized by dopamine deficiency caused by nigrostriatal dopaminergic nerve cell apoptosis. With the continued aging of the population, the incidence of PD increases yearly^1^. As the pathogenesis remains obscure, therapeutic options of PD are mainly symptomatic therapies and levodopa (L-DOPA) remains the most effective drug since the 1960s^2^. However, long-term administration of L-DOPA has limited clinical applications due to the adverse side effects with long-term use^3^. Therefore, the molecular mechanism of nigrostriatal dopaminergic nerve cell apoptosis needs to be elucidated and is of great significance for improving therapeutic strategies for the treatment of PD.

Studies have found a close link between mitochondrial dysfunction and PD pathogenesis^4-6^. Mitochondria participate in the regulation of cellular physiological functions, including cellular homeostasis, cell growth, division, and energy metabolism, specifically as it relates to apoptosis^7^. Mitochondrial dysfunction is critical to PD pathogenesis, and restoration of mitochondrial function may reduce dopaminergic nerve cell apoptosis, thereby attenuating dopamine failure and improving PD symptoms^8^. Moreover, mitochondria are dynamic and undergo frequent fission and fusion regulated by a variety of dynamic proteins, such as dynamic-related protein 1 (Drp1), fission protein 1 (Fis1), and mitofission factor (Mff) for fission and optic atrophy l (Opal) and mitofusin (Mfn) for fusion. Recent studies have shown that Drp1-induced aberrant mitochondrial fission plays an important role in the dopaminergic nerve cell apoptosis of PD. Enhanced Drp1 promotes mitochondrial fission and PD dopaminergic nerve cell apoptosis, whereas inhibited Drp1 reverses aberrant mitochondrial fission, reduces nerve cell apoptosis and improves PD symptoms^5,9-12^. Drp1 is a GTPase; once Drp1 is activated, Drp1 translocates from the cytosol to the outer mitochondrial membrane (i.e., mitochondrial translocation), forms a ring structure around the mitochondria and changes the distance and angle of molecules, gradually compressing the mitochondria until they are fractured by GTP hydrolysis, resulting in fission of mitochondria followed by cytochrome c (Cyto C) release and caspase activation, and eventually leading to apoptosis^13-16^. However, the upstream regulatory mechanism of Drp1-mediated mitochondrial fission in PD has not yet been explored.

Rho-associated coiled-coil protein kinase 1 (ROCK1) is a member of the Ras protein family with a molecular weight of 160 kDa, and plays important regulatory role in cancer cell growth and survival, as well as invasion and metastasis of neoplasm^17^. In the field of cancer research, ROCK1 has been reported to be cleaved into activated ROCK1 with a molecular weight of 130 kDa through proteolytic cleavage of its C-terminal auto-inhibitory domain, which eventually leads to apoptosis^18^. Importantly, activated ROCK1 has been found to play an important role in regulating mitochondrial fission via dephosphorylation/activation of Drp1 in human breast cancer cells^19^. Additionally, there are also reports in the central nervous system that the specific ROCK1 activation inhibitor Y-27632 decreases dopaminergic nerve cell death in mice and primary neuron-glia cultures^20,21^, but the mechanisms remain elusive. Based on the above, we propose that ROCK1 may be involved in the pathogenesis of PD as an important upstream regulator of Drp1.

In the present study, we confirm that Drp1-mediated aberrant mitochondrial fission participates in the pathogenesis of PD. Furthermore, we evaluated the role of ROCK1 in regulating dopaminergic nerve cell apoptosis in PD. We found that ROCK1 is activated in PD, and ROCK1 knockdown or pretreatment with the ROCK1 activation inhibitor Y-27632 inhibits Drp1-mediated aberrant mitochondrial fission and dopaminergic nerve cell apoptosis *in vitro* and *in vivo*, as well as significantly improves PD symptoms in a mouse model. Our mechanistic studies revealed that activated ROCK1 promotes dopaminergic nerve cell apoptosis through dephosphorylation/activation of Drp1, resulting in aberrant mitochondrial fission, and eventually leading to PD. Furthermore, we identified the therapeutic effect of Y-27632 on a PD mouse model by suppressing Drp1-mediated aberrant mitochondrial fission and dopaminergic nerve cell apoptosis. Our study contributes to a novel insight into the pathogenesis of PD involving dopaminergic nerve cell apoptosis, and provides a mechanistic basis for the promotion of ROCK1 activation inhibitor applying in the treatment of PD.

## Results

### MPP^+^ inhibits dopamine release in PD cells

Degeneration of nigrostriatal dopaminergic nerve cells in PD can be modeled by the administration of the neurotoxin 1-methyl-4-phenylpyridinium ion (MPP^+^) *in vitro^12^*. In this study, we used MPP^+^-treated dopaminergic neuron PC12 cells as a model of PD *in vitro*. First, we evaluated the effects of MPP^+^-induced dopamine loss in PC12 cells using ELISAs. As shown in Fig. 1, exposure of PC12 cells to MPP^+^ resulted in a significant decrease in dopamine production in a dose-dependent manner. This result confirms that the *in vitro* model of PD was successfully established.

**Fig. 1.**
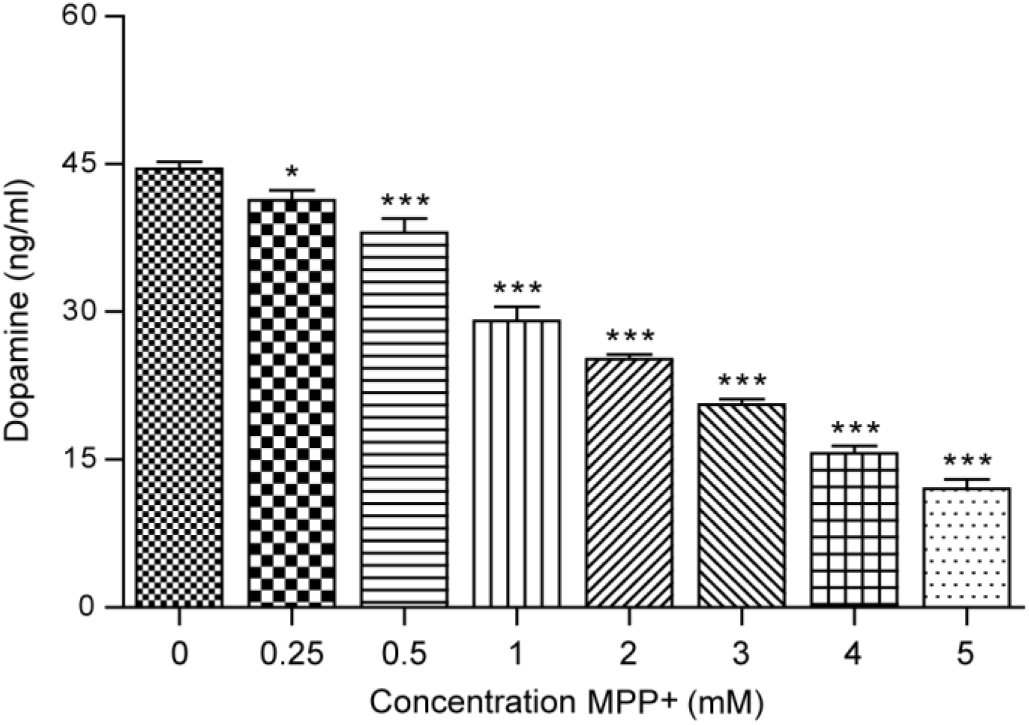
MPP^+^ inhibits dopamine release in PC12 cells. PC12 cells were treated with MPP^+^ (0, 0.25, 0.5, 1, 2, 3, 4 and 5 mM) for 24 h. The release levels of dopamine were measured using ELISA. The data are expressed as the mean ± S.D. (n = 3). \**P* < 0.05, \*\*\**P* < 0.001 vs. the control group.

### MPP^+^ induces mitochondria-dependent apoptosis in PC12 cells

To further explore the pathogenesis in the MPP^+^-induced PD model, we first studied the effect of MPP^+^ on cell viability as measured by the MTT assay. PC12 cells were treated with MPP^+^ at different concentrations and different time intervals. Our results showed that MPP^+^ resulted in significant decreases in cell viability of PC12 cells in dose- and time-dependent manners (Fig. 2a, b).

**Fig. 2.**
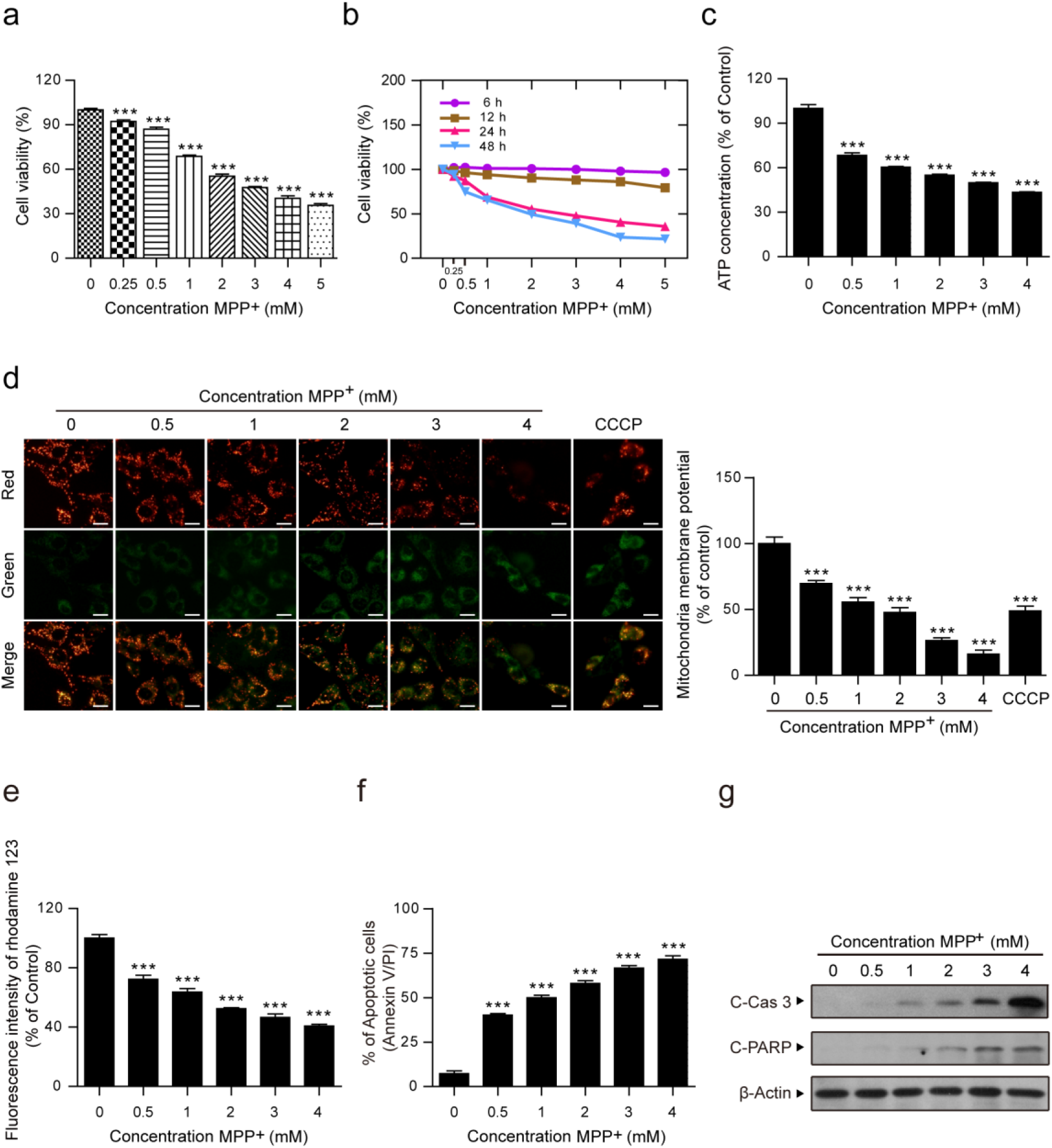
MPP^+^ induces mitochondria-dependent apoptosis in PC12 cells. PC12 cells were treated with various concentrations of MPP^+^ (0, 0.25, 0.5, 1, 2, 3, 4 and 5 mM) for 24 h (**a**) or at different time intervals (**b**), and the viability of PC12 cells was measured by MTT assays. **c** PC12 cells were treated with MPP^+^ (0, 0.5, 1, 2, 3, and 4 mM) for 24 h and the concentrations of ATP were determined using an ATP Determination Kit. **d** Mitochondrial membrane potential was measured by JC-1 staining. CCCP (10 μM) was used as the positive control. Scale bars: 20 μm. The fluorescence intensity ratio of JC-1 aggregates (red) to JC-1 monomers (green) represents the mitochondrial membrane potential. **e** Rhodamine 123 fluorescence intensity was detected by microplate reader. **f** The apoptosis cell rate was measured by flow cytometry using Annexin V-FITC/PI staining. **g** The expression of Cleaved Caspase 3 (C-Cas 3) and Cleaved PARP (C-PARP) in whole-cell lysates was determined by western blot analysis. The data are expressed as the means ± S.D. (n = 3). *\*\*\*P* < 0.001 vs. the control group.

ATP, as the most important energy molecule, plays a crucial role in the cellular physiological and pathogenic processes. ATP depletion is always an indicator of mitochondrial dysfunction^22-24^. As shown in Fig. 2c, the content of ATP rapidly decreased in the MPP^+^-treated cells in a dose-dependent manner. The loss of mitochondrial membrane potential is also another indicator of mitochondrial dysfunction^25,26^. Therefore, we examined the mitochondrial membrane potential using JC-1 and rhodamine 123 staining. The mitochondrial membrane potential of the cells using JC-1 staining is represented by the ratio of JC-1 aggregates (red) to JC-1 monomers (green) fluorescence intensities. CCCP was used as a positive control. Our results show that MPP^+^ dose-dependently decreased red fluorescence intensities and increased green fluorescence intensities and that the ratio of red to green fluorescence intensities decreased significantly (Fig. 3d). Rhodamine 123, which is specifically located on mitochondria, is also widely used to detect mitochondrial membrane potential based on fluorescence intensity^27^. Our results show that cells treated with MPP^+^ caused dose-dependent decreases in the fluorescence intensity of rhodamine 123 (Fig. 2e). Collectively, both the decrease of ATP concentration and depletion of mitochondrial membrane potential suggest that MPP^+^ induces mitochondrial dysfunction in PC12 cells.

**Fig. 3.**
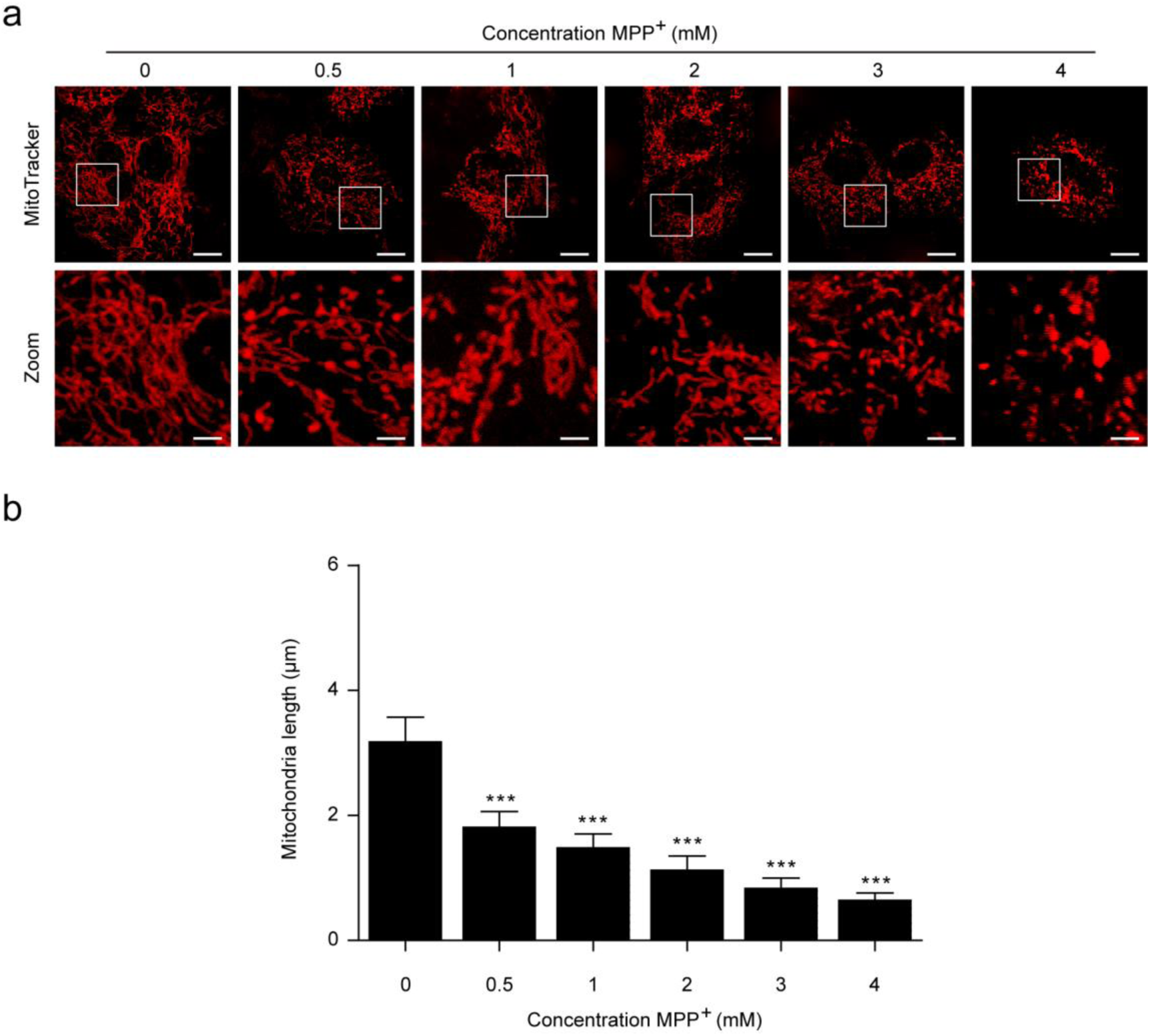
MPP^+^ induces aberrant mitochondrial fission in PC12 cells. **a** Cells were transfected with DsRed-Mito plasmid and the mitochondria morphology was viewed by confocal microscopy. Scale bars: 10 μm. **b** Quantifications of mitochondrial length were measured by Imaris software. The data are expressed as the means ± S.D. (n = 3). \*\*\**P* < 0.001 vs. the control group.

Mitochondrial dysfunction is an important indicator of mitochondria dependent apoptosis^28-30^. To investigate whether the MPP^+^-mediated mitochondrial dysfunction result in the induction of apoptosis, we used flow cytometry (Annexin V-FITC^+^/PI^-^) to identify apoptotic cells. We found that MPP^+^ led to a dose-dependent increase in the percentage of apoptotic cells (Fig. 2f). Consistent with these findings, MPP^+^ caused cleavage/activation of classical apoptosis-related proteins, such as caspase 3 and PARP (Fig. 2g). Taken together, these findings suggest that MPP^+^ induces mitochondria-dependent apoptosis in PC12 cells

### MPP^+^ induces aberrant mitochondrial fission in PC12 cells

Increasing evidence indicates that mitochondrial fission participates in the initiation of mitochondrial apoptosis^29,30^. To exam the effects of MPP^+^ on mitochondrial fission, the DsRed-Mito plasmid was transfected into cells before MPP^+^ treatment. Confocal laser scanning microscopy studies indicated that the average length of mitochondria was remarkably decreased in MPP^+^-treated cells in a dose-dependent manner (Fig. 3a, b). These results reveal that MPP^+^ induces mitochondrial apoptosis via mitochondrial fission in PC12 cells.

### MPP^+^ induces Drp1-dependent aberrant mitochondrial fission and apoptosis

Mitochondria are dynamic organelles, which undergo frequent fission and fusion. Dynamin-related protein 1 (Drp1) is responsible for mitochondrial fission through its translocation from the cytosol to mitochondria (i.e., mitochondrial translocation)^29,31,32^. Therefore, we next investigated whether mitochondrial translocation of Drp1 is a key event in MPP^+^-induced mitochondrial fission. Exposure of PC12 cells to MPP^+^ resulted in a significant increase in the levels of Drp1 in mitochondria and decrease in the cytosol in a dose dependent manner (Fig. 4a). To further verify the critical function of Drp1 on MPP^+^-induced aberrant mitochondrial fission in a PD cell culture model, lentiviral shDrp1 was used to specifically suppress the expression of Drp1 (Fig. 4b). Confocal laser scanning microscopy demonstrated that knockdown of Drp1 significantly increased the average length of mitochondria, suggesting that Drp1 knockdown inhibited MPP^+^-induced aberrant mitochondrial fission (Fig. 4c, d). Depletion of Drp1 attenuated MPP^+^-induced ATP loss compared to transfection with shCon (Fig. 4e). Moreover, Drp1 knockdown blocked MPP^+^-induced activation of caspase 3 and PARP as well as apoptosis (Fig. 4f, g). Taken together, these findings indicate that Drp1 is required for MPP^+^-induced aberrant mitochondrial fission and apoptosis.

**Fig. 4.**
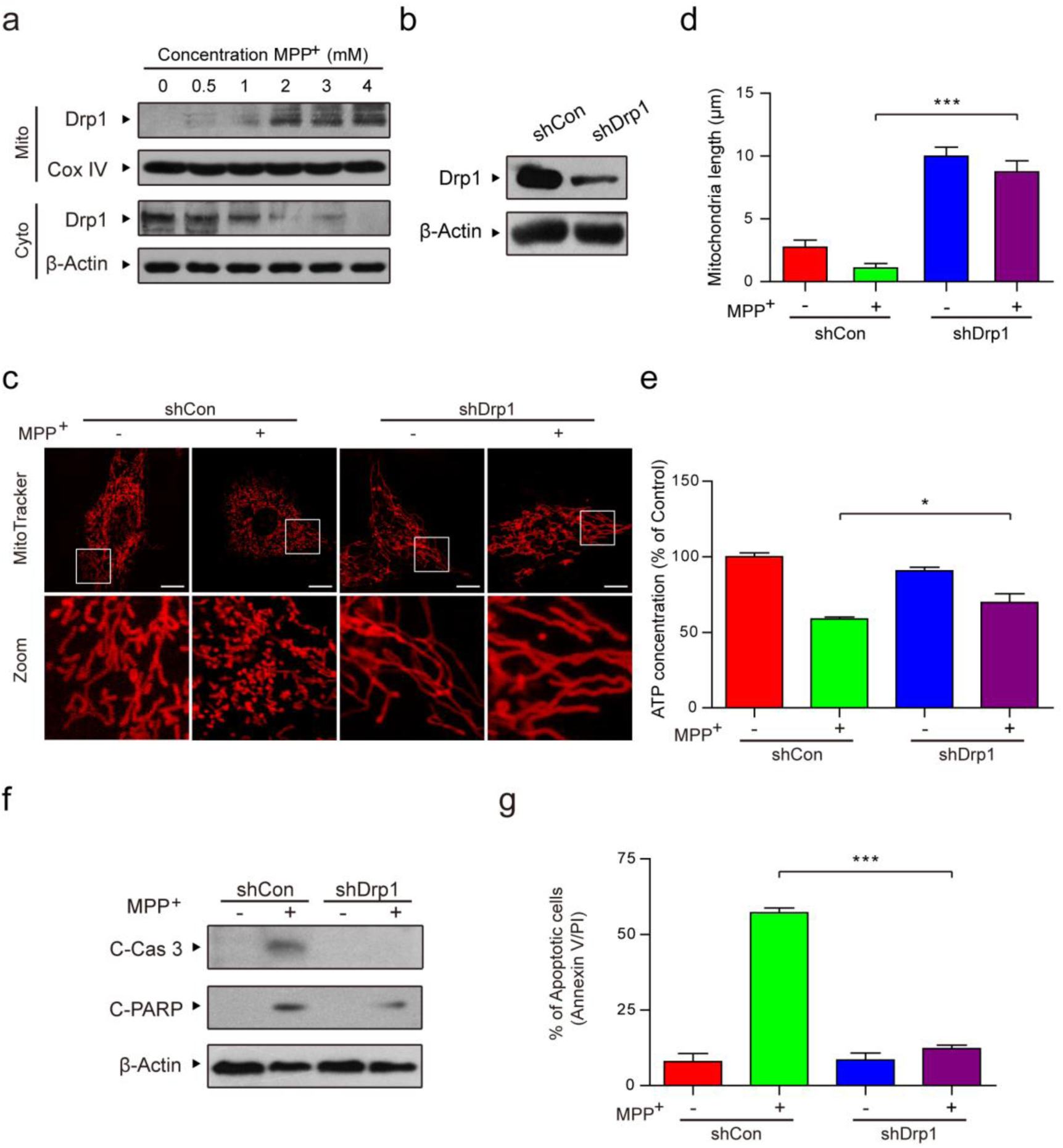
MPP^+^ induces Drp1-dependent aberrant mitochondrial fission and apoptosis. **a** PC12 cells were treated with various concentrations of MPP^+^ (0, 0.5, 1, 2, 3, and 4 mM) for 24 h. The expression of Drp1 in mitochondrial lysates (Mito) and in cytosolic fractions (Cyto) was determined by western blot analysis. **b** Stably expressed Non-Target shRNA (shCon) or Drp1 shRNA (shDrp1) PC12 cells were confirmed by western blot analysis. **c** Cells were transfected with DsRed-Mito plasmid and the mitochondria morphology was viewed by confocal microscopy. Scale bars: 10 μm. **d** Quantifications of mitochondrial length were performed using Imaris software, **e** The concentrations of ATP were determined using an ATP Determination Kit. **f** The expression of C-Cas 3 and C-PARP in whole-cell lysates was determined by western blot analysis. **g** The apoptosis cell rate was measured by flow cytometry using Annexin V-FITC/PI staining. The data are expressed as the means ± S.D. (n = 3). \**P* < 0.05, \*\*\**P* < 0.001.

### ROCK1 activation is involved in MPP^+^-induced aberrant mitochondrial fission and apoptosis through dephosphorylation/activation of Drp1

ROCK1 has been reported to play an important regulatory role in apoptosis^17,18^. Our results revealed that MPP^+^ resulted in a significant decrease in the expression of ROCK1 and increase in the expression of cleaved ROCK1 (CF: cleavage fragment) in a dose-dependent manner (Fig. 5a). ROCK1 activation is reportedly involved in the regulation of mitochondrial translocation of Drp1 and mitochondrial fission through its dephosphorylation at Ser 637 in human breast cancer cells^19^. As shown in Fig. 5b, the serine phosphorylation site is highly conserved among species and is located at the GTPase effector domain (GED) of Drp1, which suggests that Ser 656/600 in rat/mouse corresponds to Ser 637 in human due to the consensus sequence motif of ROCK substrates (R-X-X-S where R is arginine and S is serine)^33^. Thus, we identified Ser 656 in the rat Drp1 as a potential phosphorylation site for ROCK1 and next examined whether MPP^+^ had an effect on the phosphorylation state of rat Drp1 at Ser 656 in PC12 cells. A dose-dependent decrease in the level of p-Drp1 at Ser 656 (i.e., increase in dephosphorylation at Ser 656) was detected in the cells exposed to MPP^+^ (Fig. 5c). To further confirm these findings, we stably knocked down ROCK1 using a lentivirus shRNA approach (Fig. 5d). We next investigated whether ROCK1 activation was required for Drp1 translocation to mitochondria mediated by MPP^+^. ROCK1 knockdown reversed Drp1 mitochondrial translocation and dephosphorylation at Ser 656 (Fig. 5e). ROCK1 knockdown also blocked MPP^+^-mediated mitochondrial fission (Fig. 5f, g). Furthermore, knockdown of ROCK1 attenuated MPP^+^-induced ATP loss, caspase 3 and PARP activation, and apoptosis (Fig. 5h-j). Taken together, these results suggest that activated ROCK1 is involved in MPP^+^-induced aberrant mitochondrial fission and apoptosis through Drp1 dephosphorylation at Ser 656 in a PD cell culture model.

**Fig. 5.**
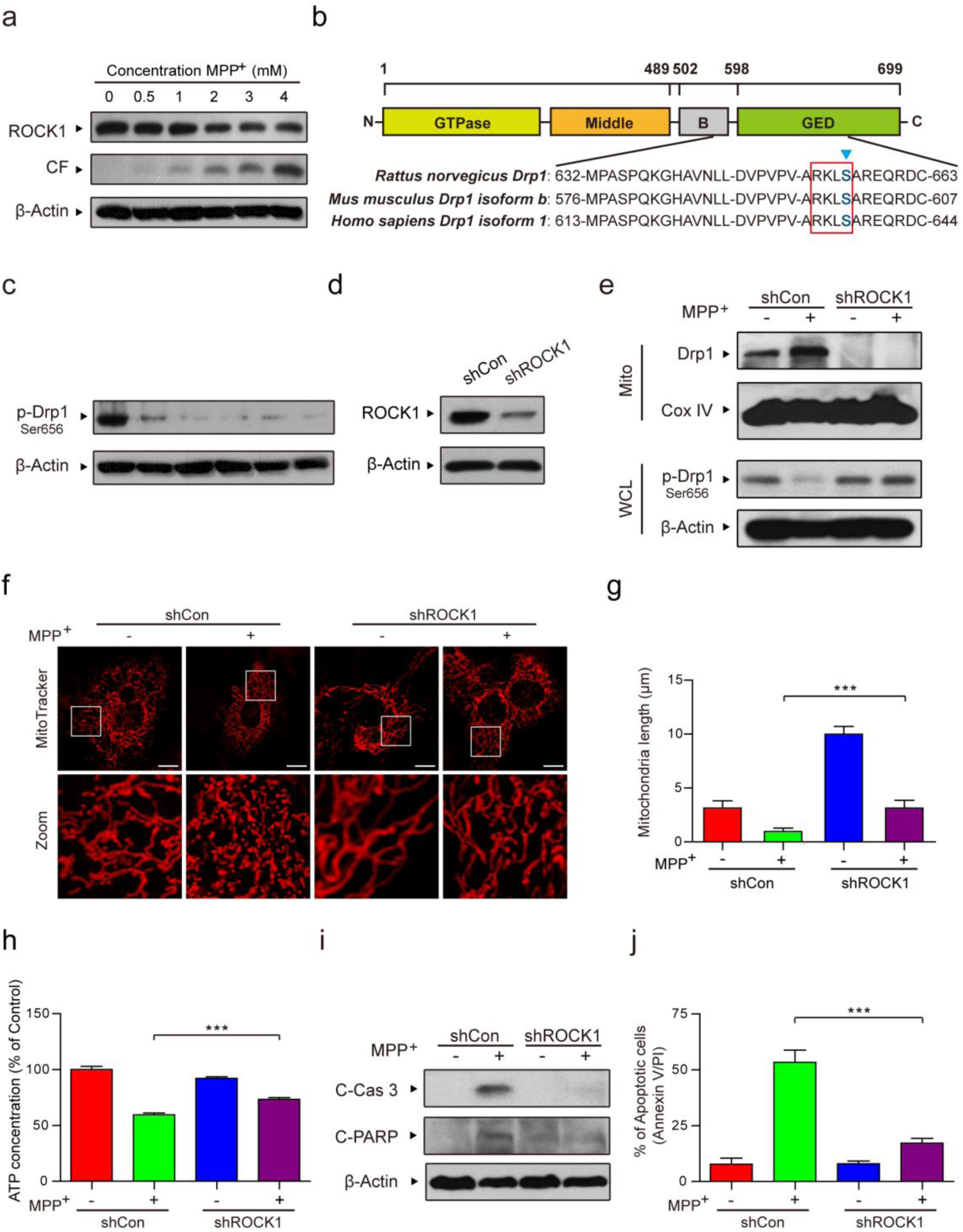
ROCK1 activation is involved in MPP^+^-induced aberrant mitochondrial fission and apoptosis through dephosphorylation/activation of Drp1. **a** PC12 cells were treated with various concentrations of MPP^+^ (0, 0.5, 1, 2, 3, and 4 mM) for 24 h. The expression of ROCK1, cleaved ROCK1 and p-Drp1 (Ser 656) in whole-cell lysates was determined by western blot analysis. CF represents the cleavage fragment of ROCK1. **b** Domain structure of rat Drp1. Sequences from several Drp1 isoforms were aligned to show the conserved motifs. **c** The expression of p-Drp1 (Ser 656) in whole-cell lysates was determined by western blot analysis. **d** Stably expressed shCon or ROCK1 shRNA (shROCK1) PC12 cells were confirmed by western blot analysis. **e** The expression of Drp1 in mitochondrial lysates (Mito) and p-Drp1 (Ser 656) in whole-cell lysates (WCL) was determined by western blot analysis. **f** Cells were transfected with DsRed-Mito plasmid and the mitochondria morphology was viewed by confocal microscopy. Scale bars: 10 μm. **g** Quantifications of mitochondrial length were performed using Imaris software, **h** The concentrations of ATP were determined using an ATP Determination Kit. **i** The expression of C-Cas 3 and C-PARP in whole-cell lysates was determined by western blot analysis. **j** The apoptosis cell rate was measured by flow cytometry using Annexin V-FITC/PI staining. The data are expressed as the means ± S.D. (n = 3). *\*\*\*P* < 0.001.

### The ROCK1 activation inhibitor Y-27632 attenuates MPP^+^-induced Drp1-dependent aberrant mitochondrial fission and apoptosis through inhibition of Drp1 dephosphorylation/activation

To further verify the critical role of the activated ROCK1 in MPP^+^-induced mitochondrial fission and apoptosis, we used Y-27632, a potent ROCK1 activation inhibitor. Preincubation of cells with Y-27632 before MPP^+^ treatment remarkably inhibited MPP^+^-induced ROCK1 activation, Drp1 dephosphorylation (Ser 656) and Drp1 mitochondrial translocation (Fig. 6a-c). Y-27632 also significantly blocked the MPP^+-^mediated mitochondrial fission (Fig. 6d, e). Furthermore, Y-27632 markedly decreased MPP^+^-induced activation of caspase 3 and PARP, as well as apoptosis (Fig. 6f, g). Collectively, our results confirmed that activated ROCK1 plays a critical role in MPP^+^-induced Drp1-dependent mitochondrial fission and apoptosis in PD cell culture models.

**Fig. 6.**
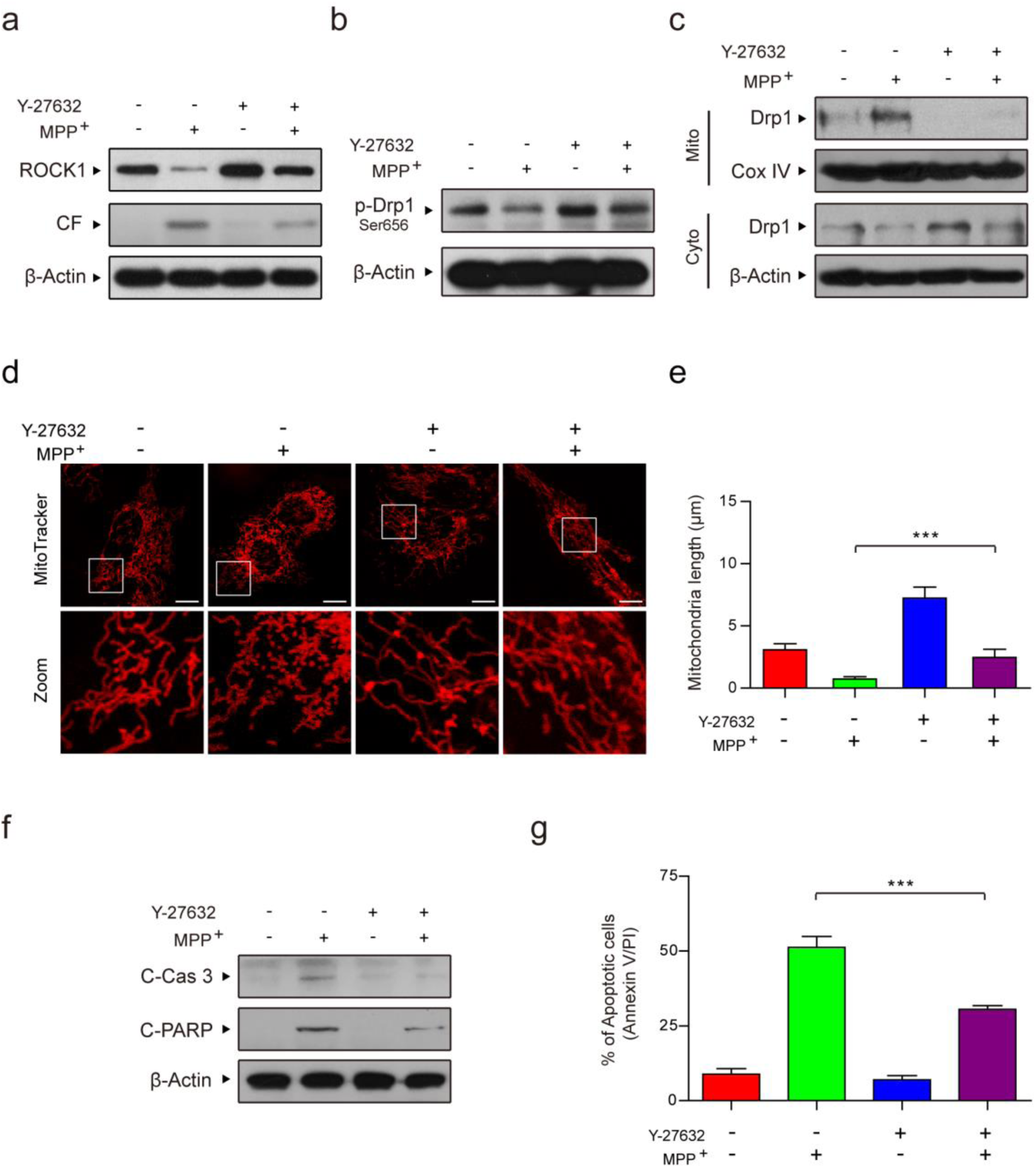
ROCK1 activation inhibitor Y-27632 attenuates MPP^+^-induced Drp1-dependent aberrant mitochondrial fission and apoptosis through inhibition of Drp1 dephosphorylation/activation. **a** PC12 cells were pretreated with ROCK1 activation inhibitor Y-27632 (50 μM) for 2 h, followed by 1 mM MPP^+^ for 24 h. The expression of ROCK1 and cleaved ROCK1 in whole-cell lysates was determined by western blot analysis. **b** The expression of p-Drp1 (Ser 656) was determined by western blot analysis. **c** The expression of Drp1 in mitochondrial lysates (Mito) and in cytosolic fractions (Cyto) was determined by western blot analysis. **d** Cells were transfected with DsRed-Mito plasmid and the mitochondria morphology was viewed by confocal microscopy. Scale bars: 10 μm. **e** Quantifications of mitochondrial length were measured by Imaris software. **f** The expression of Cleaved Caspase 3 (C-Cas 3) and Cleaved PARP (C-PARP) in whole-cell lysates was performed using by western blot analysis. **g** The apoptosis cell rate was measured by flow cytometry using Annexin V-FITC/PI staining. The data are expressed as the means ± S.D. (n = 3). \*\*\**P* < 0.001.

### The ROCK1 activation inhibitor Y-27632 improves symptoms in MPTP-induced PD mice through inhibiting Drp1-dependent aberrant mitochondrial fission and apoptosis

To verify whether our *in vitro* findings would be operative *in vivo*, we injected C57BL/6 mice with 1-methyl-4-phenyl-1, 2, 3, 6-tetrahydropyridine (MPTP, 30 mg/kg/day, intraperitoneally (i.p.)) for five consecutive days to model PD in mice. The mice in the Y-27632+MPTP group were injected with the specific ROCK1 inhibitor Y-27632 (5 mg/kg/day, i.p.) 30 min before MPTP treatment. Y-27632 remarkably inhibited MPTP-induced cleavage/activation of ROCK1 both in the substantia nigra pars compacta (SNpc) and striatum of mice (Fig. 7a). As shown in Fig. 7b, the latency of MPTP-induced PD mice to fall from the rotarod was significantly decreased, but pretreatment with Y-27632 before MPTP treatment rescued this decrease. Immunohistochemical analysis indicated that MPTP treatment resulted in a significant decrease in the number of tyrosine hydroxylase (TH, as a marker for dopamine nerve cell)-positive cells, whereas Y-27632 reversed these changes (Fig. 7c, d). The results of TH expression detected by western blot analysis were consistent with that of immunohistochemical staining (Fig. 7e). All of these findings suggest that our MPTP-induced PD mouse model was successfully established and that inhibition of ROCK1 activation using Y-27632 can protect dopamine nerve cells from the MPTP-mediated dopamine depletion in this *in vivo* model.

**Fig. 7.**
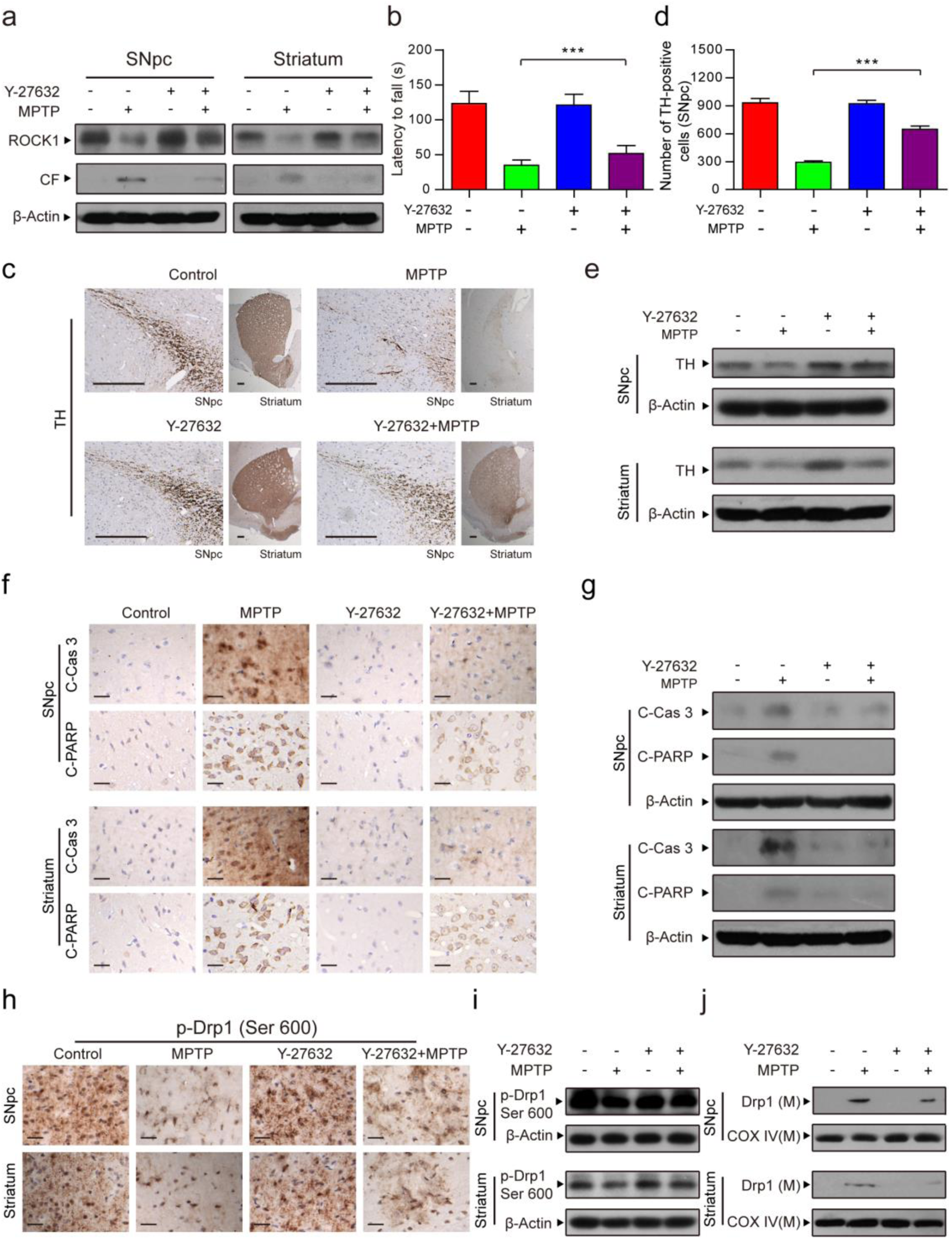
ROCK1 activation inhibitor Y-27632 improves symptoms of MPTP-induced PD mouse through inhibiting Drp1-dependent aberrant mitochondrial fission and dopaminergic nerve cell apoptosis. **a** The substantia nigra pars compacta (SNpc) of the midbrain and the striatum were prepared and subjected to detect the expression of ROCK1 and cleaved ROCK1 using western blot analysis. **b** The latency (time) to fall from the rotarod was recorded. **c** SNpc and striatum from each group were fixed, dehydrated and subjected to tyrosine hydroxylase (TH, as a marker for dopamine nerve cell) staining for immunohistochemical analysis. Scale bars: 200 μm. **d** The number of TH-positive dopaminergic nerve cells was measured by Adobe Photoshop CC. **e** The expression of TH in SNpc and striatum was determined by western blot analysis. **f** Immunohistochemistry staining of C-Cas 3 and C-PARP of SNpc and striatum are showed. Scale bars: 20 μm. **g** The expression of C-Cas 3 and C-PARP was determined by western blot analysis. **h** SNpc and striatum from each group were subjected to p-Drp1 (Ser 600) staining for immunohistochemical analysis. Scale bars: 20 μm. **i** The expression of p-Drp1 (Ser 600) was also determined by western blot analysis. **j** The expression of Drp1 in mitochondrial lysates (M) was determined by western blot analysis. The data are expressed as the means ± S.D. (n = 3). \*\*\**P* < 0.001.

We next examined the mechanism underlying PD *in vivo*. Immunohistochemical and western blot analysis showed that injection with Y-27632 before MPTP significantly inhibited the MPTP-mediated activation of caspase 3 and PARP (Fig. 7f, g). We also demonstrated that Y-27632 significantly decreased MPTP-mediated dephosphorylation of Drp1 (Ser 600) in the mouse (corresponding to Ser 637 in human, Fig. 5B) (Fig. 7h). Similarly, western blot analysis was also used to further confirm that Y-27632 attenuated MPTP-induced Drp1 (Ser 600) dephosphorylation and subsequently its mitochondrial translocation (Fig. 7i, j). Taken together, our findings indicate neuroprotective effects of an inhibitor of ROCK1 activation on an MPTP-induced mouse model of PD through inhibition of Drp1-dependent aberrant mitochondrial fission and apoptosis, suggesting ROCK1 and Drp1 may be potential therapeutic target for PD.

## Discussion

In the present study, we demonstrated for the first time that ROCK1 promotes dopaminergic nerve cell apoptosis through activating Drp1-mediated aberrant mitochondrial fission *in vitro* and *in vivo*. We also confirmed that the ROCK1 activation inhibitor Y-27632 has a therapeutic effect on a PD mouse model by suppressing Drp1-mediated aberrant mitochondrial fission and dopaminergic nerve cell apoptosis. Our findings provide a mechanistic basis for the promotion of ROCK1 activation inhibitor applying in the treatment of PD.

Currently, neurotoxic models are broadly used as models of PD^34^. The dopaminergic neurotoxin MPTP (active metabolite: MPP^+^) originates from discoveries in the early 1980s and has been used extensively to generate animal models of PD (Davis et al. 1979; Langston et al. 19 8 3 ^35,36^. MPTP contributes to the etiopathogenesis of PD by inducing mitochondria-targeted injury, decreasing dopamine levels, TH activity, and eliciting dopaminergic nerve cell apoptosis^12^. Given the parallels with PD, in this study, we examined the molecular mechanisms underlying dopaminergic nerve cell apoptosis using MPP^+^ and MPTP-induced cell and animal models of PD *in vitro* and *in vivo*, respectively^37^. Increasing evidence indicates that mitochondrial protein Drp1 is required for mitochondrial fission and MPP^+^-induced neurotoxicity^12,29,32^. Our results also demonstrate that knockdown of Drp1 significantly inhibited MPP^+^-induced aberrant mitochondrial fission and nerve cell apoptosis. Once Drp1 is activated, it translocates from the cytosol to the outer mitochondrial membrane and forms a ring structure around the mitochondria, resulting in fission of mitochondria followed by Cyto C release and caspase activation, eventually leading to apoptosis^38^. Additionally, dephosphorylation/activation of Drp1 at Ser 637 in human has been showed to promote its translocation from the cytosol to mitochondria and mitochondrial fission^19,29,39,40^. Consistent with these reports, our data revealed that dephosphorylated Drp1 at Ser 656/600 (rat/mouse) (corresponding to Ser 637 in human Drp1 isoform 1) increased mitochondrial translocation of Drp1 and leads to mitochondrial fission and nerve cell apoptosis in both *in vitro* and *in vivo* models of PD.

ROCK1 plays a central role in the regulation of cell adhesion, migration, proliferation and apoptosis^41^. ROCK1 is highly expressed in a variety of cancer tissues^42^, and plays an important role in the regulation of apoptosis in various types of cancer cells^43^. In human breast cancer cells, Drp1 has been reported to be an important ROCK1 substrate, and the dephosphorylation of Drp1 induced by ROCK1 stimulates its mitochondrial fission activity^19^. Moreover, the specific ROCK1 activation inhibitor Y-27632 has also been reported to be able to inhibit dopaminergic nerve cell death in the PD substantia nigra^21^. According to our findings, Drp1-mediated aberrant mitochondrial fission is more likely to act downstream of ROCK1 during dopaminergic nerve cell apoptosis in PD based on the following evidences. First, activation/cleavage of ROCK1 and dephosphorylation/activation of Drp1 were found in our PD models. Second, knockdown of ROCK1 remarkably decreased MPP^+^-induced dephosphorylation of Drp1, mitochondrial translocation of Drp1, aberrant mitochondrial fission and nerve cell apoptosis. Third, the ROCK1 activation inhibitor Y-27632 significantly improved symptoms of PD mice through inhibition of Drp1-mediated aberrant mitochondrial fission and dopaminergic nerve cell apoptosis. Taken together, our study reveals that ROCK1 plays a crucial role in the regulation of dopaminergic nerve cell apoptosis of PD via dephosphorylation/activation of Drp1-mediated aberrant mitochondrial fission.

In summary, the present findings indicate an important molecular mechanism of PD pathogenesis involving ROCK1-regulated dopaminergic nerve cell apoptosis. Importantly, a mechanism is also proposed for the first time by which ROCK1 cleavage/activation activates downstream Drp1 by dephosphorylation of Drp1 and subsequently induces aberrant mitochondrial fission, eventually resulting in nigrostriatal dopaminergic nerve cell apoptosis and decreasing dopamine release. Collectively, our findings contribute to a better understanding of PD pathogenesis and also provide a mechanistic basis for the promotion of ROCK1 activation inhibitor applying in the treatment of PD.

## Materials and Methods

### Reagents

1-methyl-4-phenyl-1, 2, 3, 6-tetrahydropyridine hydrochloride (MPTP-HCl, M0896) and 1-methyl-4-phenylpyridinium iodide (MPP^+^I^−^, D048) were purchased from Sigma-Aldrich Co. (St. Louis, MO, USA). Y-27632 (sc-216067) was obtained from Santa Cruz Biotechnology (Santa Cruz, CA).

### Cell lines and cell culture

PC12 cells were provided by the American Type Culture Collection (ATCC, Manassas, VA, USA). PC12 cells were cultured in RPMI-1640 medium supplemented with 10% (v/v) fetal bovine serum (FBS, Gibco, 10100) at 37 °C with 5% CO2 and 95% air in a humidified atmosphere.

### Plasmid constructs and lentiviral gene transfer

Drp1 shRNA (target sequence: 5'-CCGGGCTACTTTACTCCAACTTATTCTCGAGAATAAGTTGGAGTAAAGTAGCTTTTT-3') and ROCK1 shRNA (target sequence: 5'-CCGGCGGTTAGAACAAGAGGTAAATCTCGAGATTTACCTCTTGTTCTAACCGTTTTT-3') were subcloned into the pLKO.1 plasmid to construct shDrp1 and shROCK1 plasmid, respectively. Control shRNA plasmid (pLKO.1-puro plasmid, sc-108060) was purchased from Santa Cruz Biotechnology. Lentiviral packaging vectors pLP1, pLP2 and VSVG (Invitrogen, K4975) along with shDrp1 or shROCK1 plasmid were co-transfected into 293FT cells using Lipofectamine 3000 (Invitrogen, L3000015) according to the manufacturer’s instructions. After 48 h, lentivirus supernatant was harvested and transfected into the PC12 cells. The cells with stably knockdown of Drp1 or ROCK1 were subsequently selected with 5 μg/ml puromycin (Sigma, P9620).

### Dopamine detection

The cell culture supernatants treated with 1-methyl-4-phenylpyridinium ion (MPP^+^) were carefully collected after centrifuging at 3000 rpm for 20 min. The dopamine concentrations were quantified using enzyme-linked immunosorbent assays (ELISA) following the manufacturer’s instructions (Wuhan Colorful Gene Biological Technology, Wuhan, China).

### MTT assay

An MTT assay was performed to determine the effects of MPP^+^ on PC12 cell viability. Briefly, cells were seeded in 96-well plates and treated with MPP^+^, and the MTT solutions (5 mg/ml, 3-[4,5-dimethylthiazol-2-yl]-2,5-diphenyltetrazolium bromide, Sigma, USA) were added and incubated for 4 h. Absorption was measured by microplate reader (Thermo, Varioskan Flash) at 570 nm. The cell viabilities were normalized to the control group (100%).

#### Mitochondrial membrane potential assay by JC-1 and rhodamine 123 staining

The JC-1 kit (Beyotime Company, C2006) was used to measure the mitochondrial membrane potential according to the manufacturer’s instructions. Briefly, the cells were seeded in 24-well plate. After treatment with MPP^+^, cells were incubated with 1×JC-1 reagent solution for 15 min at 37 °C and washed twice with ice-cold 1×assay buffer. The cells incubated with carbonyl cyanide m-chlorophenylhydrazone (protonophore, CCCP, 10 μM) were used as the positive control. The fluorescence was observed by fluorescence microscopy (BX63, Olympus, Japan) and fluorescence intensity was calculated by ImageJ software (National Institutes of Health, USA). The fluorescence ratio of JC-1 aggregates (red) to JC-1 monomers (green) represents the mitochondrial membrane potential. The mitochondrial membrane potential was normalized to that of the control group (100%).

We also detected the mitochondrial membrane potential using rhodamine 123 staining. Briefly, following MPP^+^ treatment, cells were harvested and stained with 1 μM of rhodamine 123 in a 5% CO2 incubator for 30 min at 37 °C in the dark. Subsequently, the cells were washed twice with ice-cold PBS. The fluorescence intensity was measured by microplate reader (Thermo, Varioskan Flash) at 507 nm of excitation wavelength and 529 nm of emission. Rhodamine 123 fluorescence was normalized to that in the control group (100%).

#### Adenosine triphosphate (ATP) luminescence detection

The firefly luciferase-based ATP Determination Kit (Beyotime Company, S2006) was used to measure ATP levels according to the manufacturer’s instructions. Cells treated with various concentrations of MPP^+^ were lysed and centrifuged, and ATP detection working solution was added to the supernatant. The luminescence value is an index of the ATP level by using a microplate reader (Thermo, Varioskan Flash). The ATP level was normalized to the control group (100%).

#### Determination of apoptosis by flow cytometry

Cells were harvested by trypsin digestion and centrifuged for washing with PBS twice. Subsequently, the cells were resuspended in 100 μl 1×binding buffer mixed with 5 μL Annexin V-FITC and 10 μl propidium iodide (PI) (BD Biosciences, 556547) and incubated for 15 min at 25 °C in the dark. The apoptosis cell rate was analyzed by flow cytometry (FACScan, Beckman MoFlo XDP).

#### Western blot analysis

Cells and tissues were lysed with cell lysis buffer containing 1 mM PMSF. Mitochondrial and cytosolic fractions were extracted using the Cell Mitochondrial Isolation Kit (Beyotime Company, C3601). The concentrations of protein lysates were determined by BCA Protein Assay Kit (Beyotime Company, P0009). Then, 15-100 μg of sample protein was separated using SDS-PAGE and transferred to PVDF membranes. The membranes were blocked with 5% fat-free dry milk and then incubated with primary antibodies overnight at 4 °C. The protein bands were then incubated with horseradish peroxidase (HRP)-conjugated goat anti-rabbit (KPL, 074-1516) or goat anti-mouse (KPL, 074-1802) secondary antibody for 2 h at 25 °C and subsequently visualized by enhanced chemiluminescence reagent (Bio-Rad, 170-5061).

#### Immunofluorescence

Cells were plated on coverslips and then transfected with DsRed-Mito plasmid (Clontech Laboratories, Inc., PT3633-5) for 48 h using Lipofectamine 3000 (Invitrogen, L3000015). After treatment with MPP^+^, cells were fixed with 4% paraformaldehyde for 15 min and the mitochondria morphology was viewed under a LSM780 confocal laser scanning microscope (Zeiss, Germany). Mitochondrial length of at least randomly selected 10 cells were measured using Imaris software (version: 7.4.2) (Bitplane, Zurich, Switzerland).

#### Animals and treatment

All animal experiments were conducted with an approval from the Animal Care and Use Committee of Army Medical University. MPTP was used to establish a PD mouse model^44,45^. Male 8-week C57BL/6 mice (20-25 g) were randomly divided into 4 groups: control, MPTP, Y-27632, or Y-27632+MPTP (8 mice per group). The MPTP group and Y-27632 group were intraperitoneally (i.p.) injected with MPTP at a dose of 30 mg/kg/day and Y-27632 at a dose of 5 mg/kg/day once a day for 5 consecutive days, respectively^46^. The Y-27632+MPTP group mice, 30 min after injection of Y-27632 (5 mg/kg/day) 30 min, were injected with a dose of MPTP (30 mg/kg/day). The mice in the control group were injected with an equal volume of vehicle on the same schedule. On the 7th day after the last injection of MPTP, the mice were anesthetized with chloral hydrate (0.4 ml/100 g, i.p.). The mice were transcardially perfused with saline, followed by 4% paraformaldehyde. The brain was removed, immersion-fixed in 4% paraformaldehyde overnight, and dehydrated for 48 h with 30 % sucrose solution at 4 °C. The dehydrated brain tissues were coronally sectioned encompassing the entire substantia nigra pars compacta (SNpc) of the midbrain and striatum for immunofluorescence and immunohistochemical analysis. For western blot analysis, mice were euthanized under anesthesia with chloral hydrate, and brain tissues were quickly removed. SNpc of midbrain and striatum were dissected on ice.

#### Rotarod test

During the test, mice were placed on the rotarod (IITC Life Science, Series 8). Mice were pretrained for 3 days prior to the test. The training consisted of three consecutive runs gradually increasing from 5 rpm for 30 s up to a maximum 40 rpm in 5 min^45^. Each trial continued until the mice were unable to remain on the rod without falling for up to 120 s^45^. The latency (time) until the mice fell from the rotarod was recorded and the average time of three tests was analyzed for statistical analyses.

#### Statistical analysis

Data are expressed as the mean ±S.D. from at least three independent experiments. The statistical analysis was performed using one-way analysis of variance (ANOVA) with Dunnett test or Tukey by GraphPad Prism 5.0 statistical analysis software. The significance of differences between two groups was evaluated using t-tests.\**P*<0.05, \*\**P*< 0.01 or \*\*\**P*< 0.001 were regarded as a statistically significant difference.

#### Data availability

The original immunoblots gels are provided as Supplementary Figs 1–6. The authors declare that all the data supporting the findings of this study are available within the article (or the Supplementary Information) from the corresponding author on reasonable request.

## Acknowledgments

This work was supported by the National Natural Science Foundation of China (Grant No. 31600806 and 81703481), Chongqing Natural Science Foundation Program (Grant No. cstc2015shmszx120078) and Clinical Research Projects of second Affiliated Hospital, Army Medical University (Grant No. 2016YLC12).

## Author contributions

Q.Z., C.H., J.H., G.L. and R.Z. designed experiments; Q.Z., C.H., J.H., W.L., W.L., F.L., Q.T. and YL. performed all experiments; Q.Z., C.H., J.H., Q.W., M.Z. and F.S. analyzed the data; Q.Z., G.L. and R.Z. wrote the paper.

## Abbreviations

ATP: Adenosine triphosphate
C-Cas 3: Cleaved Caspase 3
CCCP: carbonyl cyanide m-chlorophenylhydrazone
COX IV: cytochrome c oxidase subunit IV isoform 1
C-PARP: Cleaved PARP
Cyto: cytosolic fractions
Cyto C: cytochrome c
Drp1: dynamin-related protein 1
Fis1: fission protein 1
Mito: mitochondrial lysates
i.p.: intraperitoneally
L-DOPA: levodopa
Mff: mitofission factor
Mfn: mitofusin
MPP^+^: 1-methyl-4-phenylpyridinium ion
MPTP: 1-methyl-4-phenyl-1, 2, 3, 6-tetrahydropyridine
Opal: optic atrophy 1
PD: Parkinson's disease
ROCK1: Rho-associated coiled-coil protein kinase 1
SNpc: substantia nigra pars compacta
TH: tyrosine hydroxylase
WCL: whole-cell lysates.

## References

1. Zou, Y. M., Liu, J., Tian, Z. Y. et al. Systematic review of the prevalence and incidence of Parkinson’s disease in the People’s Republic of China. Neuropsychiatr. Dis. Treat. 11, 1467–1472 (2015).

2. Katzenschlager, R. & Lees, A. J. Treatment of Parkinson’s disease: levodopa as the first choice. J. Neurol. 249 Suppl 2, II19–24 (2002).

3. Cacabelos, R. Parkinson’s Disease: From Pathogenesis to Pharmacogenomics. Int J Mol Sci 18 (2017).

4. Giannoccaro, M. P., La Morgia, C., Rizzo, G. et al. Mitochondrial DNA and primary mitochondrial dysfunction in Parkinson’s disease. Mov. Disord. 32, 346–363 (2017).

5. Rappold, P. M., Cui, M., Grima, J. C. et al. Drp1 inhibition attenuates neurotoxicity and dopamine release deficits in vivo. Nature Communications 5 (2014).

6. Vives-Bauza, C., Tocilescu, M., de Vries, R. L. A. et al. Control of mitochondrial integrity in Parkinson’s disease. Recent Advances in Parkinsons Disease: Basic Research 183, 99–113 (2010).

7. Ishihara, N., Otera, H., Oka, T. et al. Regulation and Physiologic Functions of GTPases in Mitochondrial Fusion and Fission in Mammals. Antioxidants & Redox Signaling 19, 389–399 (2013).

8. Inoue, N., Ogura, S., Kasai, A. et al. Knockdown of the mitochondria-localized protein p13 protects against experimental parkinsonism. EMBO Rep (2018).

9. Wang, W. Z., Wang, X. L., Fujioka, H. et al. Parkinson’s disease-associated mutant VPS35 causes mitochondrial dysfunction by recycling DLP1 complexes. Nat. Med. 22, 54–+ (2016).

10. Burte, F., Carelli, V., Chinnery, P. F. et al. Disturbed mitochondrial dynamics and neurodegenerative disorders. Nature Reviews Neurology 11, 11–24 (2015).

11. Wang, X., Yan, M. H., Fujioka, H. et al. LRRK2 regulates mitochondrial dynamics and function through direct interaction with DLP1. Hum. Mol. Genet. 21, 1931–1944 (2012).

12. Wang, X. L., Su, B., Liu, W. H. et al. DLP1-dependent mitochondrial fragmentation mediates 1-methyl-4-phenylpyridinium toxicity in neurons: implications for Parkinson’s disease. Aging Cell 10, 807–823 (2011).

13. Wang, J. X., Li, Q. & Li, P. F. Apoptosis Repressor with Caspase Recruitment Domain Contributes to Chemotherapy Resistance by Abolishing Mitochondrial Fission Mediated by Dynamin-Related Protein-1. Cancer Res. 69, 492–500 (2009).

14. Tanaka, A. & Youle, R. J. A chemical inhibitor of DRP1 uncouples mitochondrial fission and apoptosis. Mol. Cell 29, 409–410 (2008).

15. Estaquier, J. & Arnoult, D. Inhibiting Drp1-mediated mitochondrial fission selectively prevents the release of cytochrome c during apoptosis. Cell Death Differ. 14, 1086–1094 (2007).

16. Frank, S., Gaume, B., Bergmann-Leitner, E. S. et al. The role of dynamin-related protein 1, a mediator of mitochondrial fission, in apoptosis. Dev. Cell 1, 515–525 (2001).

17. Wei, L., Surma, M., Shi, S. et al. Novel Insights into the Roles of Rho Kinase in Cancer. Arch. Immunol. Ther. Exp. (Warsz.) 64, 259–278 (2016).

18. Vemula, S., Shi, J. J., Hanneman, P. et al. ROCK1 functions as a suppressor of inflammatory cell migration by regulating PTEN phosphorylation and stability. Blood 115, 1785–1796 (2010).

19. Li, G. B., Zhou, J., Budhraja, A. et al. Mitochondrial translocation and interaction of cofilin and Drp1 are required for erucin-induced mitochondrial fission and apoptosis. Oncotarget 6, 1834–1849 (2015).

20. He, Q., Li, Y. H., Guo, S. S. et al. Inhibition of Rho-kinase by Fasudil protects dopamine neurons and attenuates inflammatory response in an intranasal lipopolysaccharide-mediated Parkinson’s model. Eur. J. Neurosci. 43, 41–52 (2016).

21. Borrajo, A., Rodriguez-Perez, A. I., Villar-Cheda, B. et al. Inhibition of the microglial response is essential for the neuroprotective effects of Rho-kinase inhibitors on MPTP-induced dopaminergic cell death. Neuropharmacology 85, 1–8 (2014).

22. Singleterry, J., Sreedhar, A. & Zhao, Y. F. Components of cancer metabolism and therapeutic interventions. Mitochondrion 17, 50–55 (2014).

23. Brandon, M., Baldi, P. & Wallace, D. C. Mitochondrial mutations in cancer. Oncogene 25, 4647–4662 (2006).

24. Skulachev, V. P. Mitochondrial physiology and pathology; concepts of programmed death of organelles, cells and organisms. Mol. Aspects Med. 20, 139–184 (1999).

25. Pokorny, J., Pokorny, J., Kobilkova, J. et al. Targeting mitochondria for cancer treatment - two types of mitochondrial dysfunction. Prague Med. Rep. 115, 104–119 (2014).

26. Safe, S. Targeting Apoptosis Pathways in Cancer-Letter. Cancer Prev Res 8, 338–338 (2015).

27. Murugan, C., Rayappan, K., Thangam, R. et al. Combinatorial nanocarrier based drug delivery approach for amalgamation of anti-tumor agents in bresat cancer cells: an improved nanomedicine strategies. Sci. Rep. 6, 34053 (2016).

28. Adams, J. M. & Cory, S. The Bcl-2 apoptotic switch in cancer development and therapy. Oncogene 26, 1324–1337 (2007).

29. Sheridan, C. & Martin, S. J. Mitochondrial fission/fusion dynamics and apoptosis. Mitochondrion 10, 640–648 (2010).

30. Perfettini, J. L., Roumier, T. & Kroemer, G. Mitochondrial fusion and fission in the control of apoptosis. Trends Cell Biol. 15, 179–183 (2005).

31. Otera, H. & Mihara, K. Molecular mechanisms and physiologic functions of mitochondrial dynamics. J. Biochem. 149, 241–251 (2011).

32. Knott, A. B., Perkins, G., Schwarzenbacher, R. et al. Mitochondrial fragmentation in neurodegeneration. Nature Reviews Neuroscience 9, 505–518 (2008).

33. Kang, J. H., Jiang, Y., Toita, R. et al. Phosphorylation of Rho-associated kinase (Rho-kinase/ROCK/ROK) substrates by protein kinases A and C. Biochimie 89, 39–47 (2007).

34. Tieu, K. A guide to neurotoxic animal models of Parkinson’s disease. Cold Spring Harb. Perspect. Med. 1, a009316 (2011).

35. Davis, G. C., Williams, A. C., Markey, S. P. et al. Chronic Parkinsonism secondary to intravenous injection of meperidine analogues. Psychiatry Res. 1, 249–254 (1979).

36. Langston, J. W., Ballard, P., Tetrud, J. W. et al. Chronic Parkinsonism in humans due to a product of meperidine-analog synthesis. Science 219, 979–980 (1983).

37. Haque, M. E., Thomas, K. J., D'Souza, C. et al. Cytoplasmic Pink1 activity protects neurons from dopaminergic neurotoxin MPTP. Proc. Natl. Acad. Sci. U. S. A. 105, 1716–1721 (2008).

38. Chang, C. R. & Blackstone, C. Dynamic regulation of mitochondrial fission through modification of the dynamin-related protein Drp1. Mitochondrial Research in Translational Medicine 1201, 34–39 (2010).

39. Cereghetti, G. M., Stangherlin, A., Martins de Brito, O. et al. Dephosphorylation by calcineurin regulates translocation of Drp1 to mitochondria. Proc. Natl. Acad. Sci. U. S. A. 105, 15803–15808 (2008)

40. Li, G. B., Fu, R. Q., Shen, H. M. et al. Polyphyllin I induces mitophagic and apoptotic cell death in human breast cancer cells by increasing mitochondrial PINK1 levels. Oncotarget 8, 10359–10374 (2017).

41. Julian, L. & Olson, M. F. Rho-associated coiled-coil containing kinases (ROCK): structure, regulation, and functions. Small GTPases 5, e29846 (2014).

42. Lochhead, P. A., Wickman, G., Mezna, M. et al. Activating ROCK1 somatic mutations in human cancer. Oncogene 29, 2591–2598 (2010).

43. Tsai, N. P., & Wei, L. N. RhoA/ROCK1 signaling regulates stress granule formation and apoptosis. Cell. Signal. 22, 668–675 (2010).

44. Santos, D. B., Colle, D., Moreira, E. L. et al. Succinobucol, a Non-Statin Hypocholesterolemic Drug, Prevents Premotor Symptoms and Nigrostriatal Neurodegeneration in an Experimental Model of Parkinson’s Disease. Mol. Neurobiol. 54, 1513–1530 (2017).

45. Guo, B., Hu, S., Zheng, C. et al. Substantial protection against MPTP-associated Parkinson’s neurotoxicity in vitro and in vivo by anti-cancer agent SU4312 via activation of MEF2D and inhibition of MAO-B. Neuropharmacology 126, 12–24 (2017).

46. Villar-Cheda, B., Dominguez-Meijide, A., Joglar, B. et al. Involvement of microglial RhoA/Rho-kinase pathway activation in the dopaminergic neuron death. Role of angiotensin via angiotensin type 1 receptors. Neurobiol. Dis. 47, 268–279 (2012).

